# RefLA: Efficient reference-guided genome assembly from long reads

**DOI:** 10.64898/2025.12.23.696147

**Authors:** XianFeng Shi, Hansen Chen, Fan Nie, Jianxin Wang

## Abstract

High-quality de novo genome assembly from long reads remains computationally demanding, while fast assemblers often compromise on contiguity or accuracy. This trade-off between speed and assembly quality presents a persistent challenge in large-scale genomic studies. Fortunately, high-quality reference genomes are now available for a growing number of species. These references can be leveraged to assist the assembly of closely related genomes, enabling efficient reconstruction without sacrificing fidelity—particularly when evolutionary divergence is moderate.We present RefLA, a reference-assisted assembler tailored for thirdgeneration long-read data. By combining alignment-guided structural refinement with localized reassembly, RefLA achieves accuracy comparable to or better than state-of-the-art de novo tools such as Flye and hifiasm, while running significantly faster. It also surpasses rapid assemblers like wtdbg2 and Shasta in both contiguity and base-level correctness.

## 1 Introduction

Genome sequencing generates vast amounts of fragmented sequences. To comprehend the complete genomic landscape, computational methods are required to piece these fragments together—a process known as genome assembly[1]. High-quality genome assemblies are fundamental for research in species evolution, gene expression, population genetics, and more. Constructing high-quality reference genomes and performing large-scale sequencing and assembly have provided novel insights into gene expression regulation, population genetics, and evolution, contributing to advancements in animal and plant breeding, speciation studies, and precision medicine[2–9].

Genome assembly can be categorized into de novo assembly and reference-guided assembly, depending on whether a reference sequence is used. De novo assembly relies on overlap relationships between reads to reconstruct the genome without a reference. A major challenge lies in genomic repeat regions, which vary in length and copy number, often leading to misassembly. Common de novo assembly strategies include those based on de Bruijn graphs (DBG), overlap-layout-consensus (OLC) graphs, and string graphs[10, 11]. Both types of algorithms generally follow these steps: first, detect overlaps between reads; second, construct an assembly graph based on these overlaps; third, simplify the graph; and finally, traverse the graph to compute the optimal path for assembly. Different sequencing data types often favor different strategies. DBG significantly simplifies computational efficiency by transforming the problem into finding Eulerian paths, making it widely used in second-generation (short-read) assemblers. However, DBG cannot fully utilize the advantages of long reads and is less suitable for high-error-rate data. In contrast, OLC graphs better accommodate long-read characteristics and tolerate a certain degree of sequencing error, making them more suitable for assembling long reads with higher error rates.

Reference-guided assembly requires a reference sequence from the target species or a closely related species. Reads are aligned to this reference to assemble the target genome. This approach avoids all-vs-all read comparisons and provides a prior information about complex repeat regions, often resulting in faster assembly. However, it can inadvertently introduce biases from the reference. To combine the high accuracy of short reads with long reads, hybrid error-correction pipelines such as PBcR[12] and Nanocorr[13] first correct long reads using short reads before performing de novo assembly. Another category, including HYBRIDSPADES[14], performs de novo assembly with short reads to produce contigs, then aligns long reads to the assembly graph to bridge, resolve repeats, and fill gaps. While hybrid correction leverages the high accuracy of short reads, it has significant limitations, such as difficulties in correcting GC-rich regions and poor performance in repeat regions[10]. Subsequently, Chen-Shan et al.[15] developed a method for assembling third-generation data without hybrid short reads, employing a hierarchical self-correction strategy with long reads, successfully applied to microbes like E.coli. The HGAP algorithm selects the longest read as a seed, aligns other shorter reads to it to generate a consensus preassembly, and then proceeds with the assembly pipeline. To scale long-read assembly to large genomes, Konstantin Berlin et al.[16] introduced the MinHash Alignment Process (MHAP) algorithm, which uses probabilistic and locality-sensitive hashing to rapidly compute read overlaps, overcoming a computational bottleneck. Integrated with the Celera Assembler, it successfully assembled low-complexity regions of CHM1. Koren et al.[17] Introduced Canu, an assembler for third-generation nanopore reads, which improved speed by using a tf-idf weighted MinHash method to find candidate overlaps and filter them based on sequence statistics, enabling differentiation of repeats with *≥* 3% divergence and improving upon the earlier Celera Assembler. Xiao et al.[18] developed MECAT, which introduced a pseudo-linear alignment scoring algorithm to filter excessive invalid alignments, reducing redundant computation and improving assembly efficiency. Ying Chen et al.[19]proposed NECAT for nanopore read assembly, featuring a two-stage error correction method to handle complex errors in nanopore data. NECAT corrects low-error-rate regions first, followed by high-errorrate regions. By avoiding direct trimming of high-error reads and employing segmental correction, NECAT improves assembly contiguity. However, these methods remain computationally expensive due to the time-consuming overlap detection and correction steps.

Mikhail et al.[20]proposed Flye, which addresses repeat assembly using an iterative strategy: first, rapidly constructing and correcting disjointigs, then building a repeat graph, aligning sequences to it, and simplifying to produce the assembly. Flye shows significant improvements over tools like Canu, especially for repeat regions. Traditional DBG is not well-suited for high-error-rate third-generation reads. Ruan J et al.[21] developed wtdbg2, which modifies the DBG approach by introducing the fuzzy Bruijn graph (FBG) concept to tolerate high error rates and using k-mer binning to improve speed. Shafin et al.[22]developed Shasta, which excels at human genome assembly. Shasta compresses homopolymers in long reads to mitigate errors, uses MinHash for overlap detection, and employs MarginPolish and HELEN for polishing to improve accuracy. With further sequencing technology advances, PacBio developed Circular Consensus Sequencing (CCS), producing high-fidelity (HiFi) reads. For PacBio HiFi data, Cheng et al.[23] developed Hifiasm, which offers faster assembly and high accuracy, though it relies on costly HiFi reads.

To leverage the prior information from high-quality reference genomes, we propose RefLA, a reference-guided assembly algorithm. RefLA fully utilizes reference information, treating it as an initial draft assembly. It comprehensively identifies divergent regions between the target and reference genomes from read alignment data, classifies these regions, and applies different correction strategies. Experiments on ONT/HiFi datasets show that RefLA achieves assembly quality comparable or superior to high-quality assemblers like Flye and Hifiasm while being significantly faster, and outperforms wtdbg2 and Shasta in quality. Its speed advantage is particularly pronounced for large genomes, where it is second only to Shasta.

## 2 Method

### 2.1 Overview of the RefLA

We present RefLA, an efficient reference-guided assembler for third-generation sequencing data. The overall design is inspired by de novo assemblers that follow an assemble-then-correct paradigm. Unlike traditional de novo assemblers, RefLA uses a reference genome as the initial assembly and then applies different correction strategies to specific regions. To mitigate bias from the reference while leveraging its information to improve quality and speed, we designed key methods for locating, classifying, and correcting divergent regions. RefLA introduces the concept of “genome divergent regions” to address assembly errors arising from genomic differences between the target and reference. A divergent region is defined as an area where alignment is abnormal due to variations (of different types and scales) between the two genomes.

The RefLA workflow is shown in Figure 1 and consists of the following main steps: reference genome selection and preprocessing, read alignment, feature extraction and candidate region identification, classification and processing of divergent regions, polishing, and scaffolding.

**Fig. 1.**
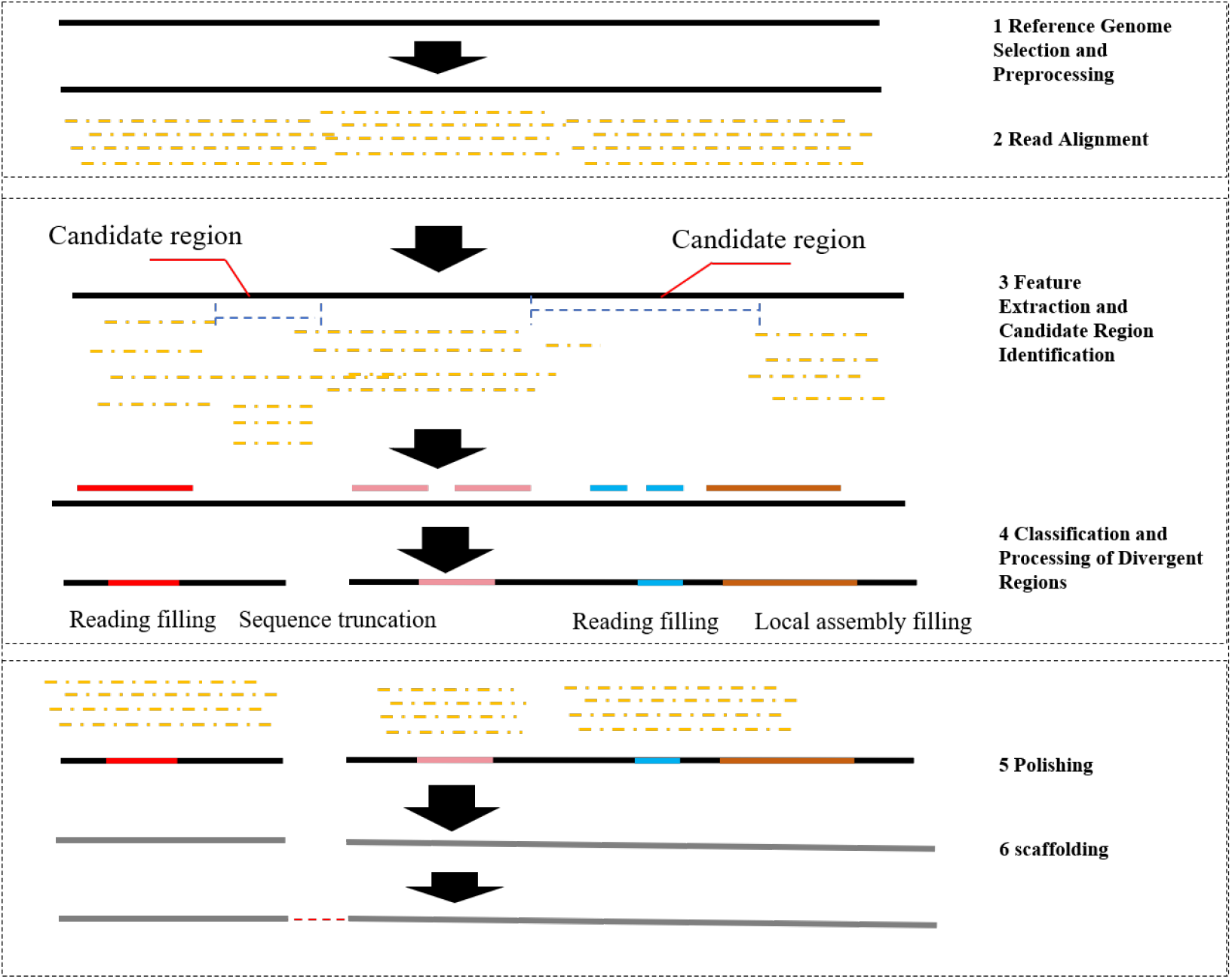
RefLA flowchart

### 2.2 Reference Genome Selection and Preprocessing

Genomes of different individuals or species typically exhibit variations. These include single nucleotide polymorphisms (SNPs) and various types of structural variations (SVs). Structural variations are generally defined as alterations *≥* 50 bp, including insertions, deletions, inversions, duplications, and translocations. SVs, involving changes of multiple bases, pose greater challenges for assembly.

Unlike general assemblers, RefLA relies on an existing reference sequence. While the reference provides prior information that can reduce assembly complexity, it also introduces bias due to genomic differences. If the divergence between the target and reference is large, resulting in low collinearity, the usable information decreases, increasing assembly difficulty. Therefore, it is recommended to select a high-quality reference genome from a closely related species. This reduces complexity and improves final assembly quality. Many reference genomes (e.g., GRCh38) contain fragmented sequences and regions filled with ‘N’s. RefLA provides an optional preprocessing step to discard fragmented contigs and retain long, continuous scaffold sequences. Based on our tests, discarding fragmented sequences from the reference is recommended.

### 2.3 Read Alignment

Minimap2, developed by Heng Li’s team, is a widely used alignment tool for mapping DNA or mRNA sequences to a reference, particularly for third-generation nanopore and HiFi reads. To reduce spurious alignments and improve speed, it employs heuristic algorithms tailored for long reads. In this study, RefLA uses Minimap2 to align long reads to the preprocessed reference genome, generating a BAM file. This file is then sorted and indexed for subsequent processing. To accelerate alignment, we adjusted default Minimap2 parameters, primarily the -w and -k parameters (e.g., -w27 -k27).

### 2.4 Feature Extraction and Candidate Region Identification

Reference-guided assembly relies on the reference genome to assemble the target. A core challenge is addressing bias introduced by differences between the reference and target sequences. We collectively term these differences “divergent regions” or “problematic regions.” Genomic differences include not only simple SVs and SNPs but also complex regions like centromeres that cannot be characterized by simple SVs. Divergent regions may arise from combinations of different variant types and sizes, significantly increasing complexity. Therefore, a highly sensitive divergence detection algorithm is needed to comprehensively identify all types of differences for subsequent correction.

We previously proposed the LRMD[24] algorithm for detecting assembly errors, which uses multiple features and effective filtering. Since detecting assembly errors is similar to detecting divergent regions, RefLA’s divergence detection algorithm adapts and improves upon LRMD’s feature extraction step. RefLA utilizes the following features, computed from the BAM file using sliding windows (partitioning the reference into windows and calculating feature values per window), for initial candidate region identification and filtering: coverage depth, read clipping, and pileup information.

### 2.5 Classification and Processing of Divergent Regions

Candidate regions from the previous step are further clustered and classified based on proximity and characteristics. According to the availability of supporting reads and the strategy for local assembly, they are categorized into: read-fill regions, assembly-fill regions, assembly-bridge regions, and assembly-extension regions. Read-filling handles regions where well-aligned reads span the divergence, allowing direct correction.

For regions not solvable by direct read filling, RefLA employs local assembly with three strategies: Assembly-filling resolves gaps where both sides belong to the same continuous block, using local assembly results spanning the gap. Assembly-bridging handles gaps where the two sides belong to different blocks (e.g., due to translocations, inversions), involving sequence truncation and re-bridging. Assembly extension further improves genome completeness. The overall classification and processing workflow is shown in Figure 2. The main categories and handling methods are as follows:

**Fig. 2.**
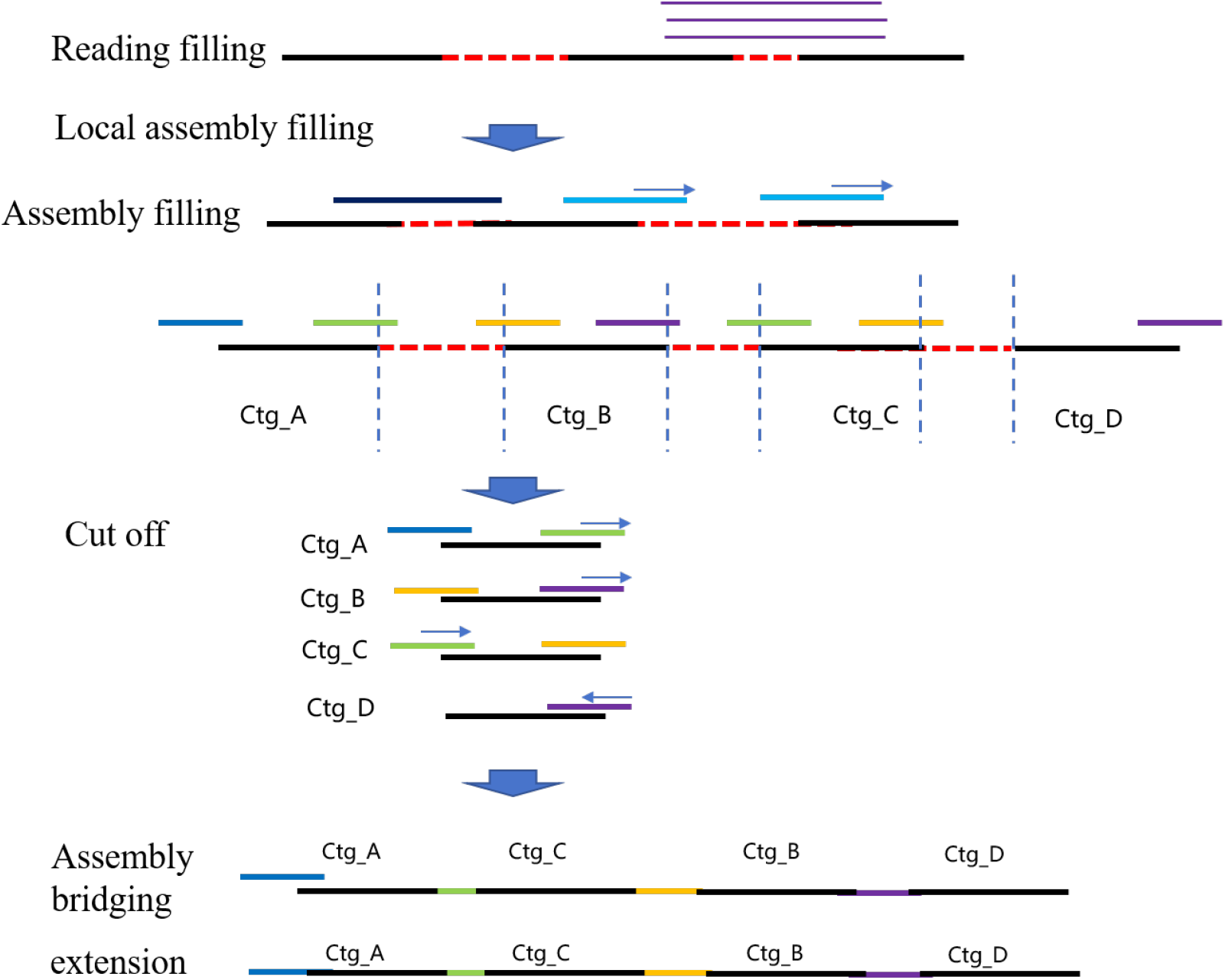
Differential Region Classification and Processing Flow

#### 2.5.1 Divergence Classification Criteria

Classification follows these rules:

1. **Read-fill Criterion:** For each clustered candidate region, check if there are sufficient supporting reads. If there are at least *N* reads supporting the region and the region’s average depth Avg_ dp *≥* 0.5 *×* whole _ dp (where whole dp is the global average depth), the region is classified as *read-fill. N* is set to 3. A supporting read must satisfy:
  a. It is a primary alignment (secondary/supplementary alignments filtered);
  b. It spans the region (read ref start *<* region start *−* 1000 and read ref end *>* region end + 1000);
  c. Mapping quality MAPQ *≥* 20, alignment length *>* 10,000, and alignment identity *≥* 0.9;
  d. Left/right clip length *<* 1000.
2. **Assembly-fill Criterion:** Regions not classified as read-fill are merged and classified for local assembly. Reads are collected from these regions, including unmapped reads and highly clipped reads. Local assembly is performed using tools such as Hifiasm (for HiFi data) or Shasta (for Nanopore data). For each resulting local assembly gap, it is classified as *assembly-fill* if either:
  a. There is an assembled contig spanning the gap; or
  b. A single contig aligns in the same orientation to both sides of the gap (split alignment).
  c. Such gaps are filled using the assembly sequence.
3. **Assembly-bridge Criterion:** Gaps not resolved by assembly-fill are collected. The reference sequence is truncated at these gaps, producing contig fragments. Bridging sequences connecting these fragments are identified. For each bridging sequence:
  a. If it connects to only one location, bridging fails.
  b. If it connects to two locations, check the alignment orientations (strand_1_, strand_2_) and the sides of the contigs it connects to (bound_1_, bound_2_). If (strand_1_ = strand_2_ and bound_1_ ≠ bound_2_),

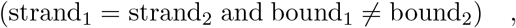

Then attempt bridging.
  c. If it connects to three locations, select the two connections with the best overlap for bridging. More than three connections lead to failure.
4. **Assembly-extension Criterion:** All bridging sequences that failed or connect to only one location are classified for *assembly-extension*. These sequences are used to extend the contig at their single connection point.

#### 2.5.2 Divergence Processing Methods

Processing follows these rules:

1. **Read-fill Processing:** As shown in Figure 3, for a region [*a, b*] on the reference, let Seq_1_ and Seq_2_ be the sequences immediately left and right. The best supporting read (based on highest alignment identity, Formula 1, where NM is edit distance from BAM, read len is original read length) is selected. Its CIGAR string is parsed to find positions *c* and *d* on the read corresponding to reference positions *a* and *b*. Let Seq be the read sequence and strand its alignment orientation. If strand is forward, filling sequence P seq = Seq[*c*:*d*]. If strand is reverse, P seq = reverse complement(Seq)[*c*:*d*]. The final corrected sequence for the region becomes Seq_1_ + P seq + Seq_2_.

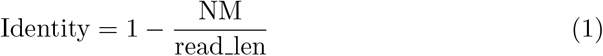
2. **Local Assembly Method:** Local assembly involves collecting three types of reads:
  a. Reads from the target gap region (extracted by read ID from original FASTQ using pysam);
  b. Unmapped reads (extracted from BAM using samtools and converted to FASTQ);
  c. Highly clipped reads.
  Highly clipped reads are identified before read-fill processing: for each candidate read-fill region, collect reads with clip length exceeding a threshold; if the number of such reads exceeds a threshold, add them to a candidate set. After read-filling, realign these candidate clipped reads to the corrected sequence; those that remain highly clipped are retained for local assembly. The combined read set is assembled using Hifiasm (HiFi data) or Shasta (Nanopore data).
3. **Assembly-fill Processing:** As shown in Figure 4, for a gap region [*a, b*], two scenarios exist.
  a. If a locally assembled contig spans the gap, processing is identical to read-fill using the contig sequence.
  b. If a single contig aligns in split-read fashion to both sides of the gap with the same orientation, parse the CIGAR to find positions *c* and *d* on the contig corresponding to reference positions *a* and *b* (*c ≤ d*). If alignment is forward, P_seq = Seq_contig[*c*:*d*]; if reverse, P seq = reverse complement(Seq contig)[*c*:*d*]. Corrected sequence: Seq_1_ +P seq+ Seq_2_.
4. **Assembly-bridge Processing:** As shown in Figure 5, for a bridging sequence Seq connecting to contigs at sites *a, b, c* … For connections to two sites (contigs contig_1_ and contig_2_), with alignment strands strand_1_, strand_2_ and connection sides bound_1_, bound_2_. Proceed only if: (strand_1_ = strand_2_ & bound_1_ ≠ bound_2_) OR (strand_1_ ≠ strand_2_ & bound_1_ = bound_2_). Let position *d* on Seq connect to site *a*, and position *e* on Seq connect to site *b*. Let Seq_1_ and Seq_2_ be the contig sequences.
  a. If strand_1_ = strand_2_ and bound_1_ is the right side of contig_1_:
    - If forward alignment, bridging sequence B seq = Seq[*d*:*e*];
    - If reverse, B seq = reverse complement(Seq)[*d*:*e*]. Final bridged sequence: Seq_1_ + B seq + Seq_2_.
  b. If strand_1_ ≠ strand_2_, bound_1_ is right side of contig_1_ (strand_1_ forward), bound_2_ is right side of contig_2_ (strand_2_ reverse): B seq = Seq[*d*:*e*]. Final bridged sequence: Seq_1_ + B seq + reverse complement(Seq_2_). Upon successful bridging, contig_1_ and contig_2_ are replaced by the new contig_3_. For sequences connecting three sites, select the two with the longest overlap for bridging. More than three connections result in failure.
  c. **Assembly-extension Processing:** As shown in Figure 6, for bridging sequences that failed or connect to only one site, use them to extend the connected contig. Suppose contig_1_ (Seq_1_) connects at its right side to position *a* on bridging sequence Seq. If Seq aligns forward, extended contig = Seq_1_ + Seq[*a*:]. If Seq aligns reverse, extended contig = Seq_1_ + reverse complement(Seq)[*a*:].

**Fig. 3.**
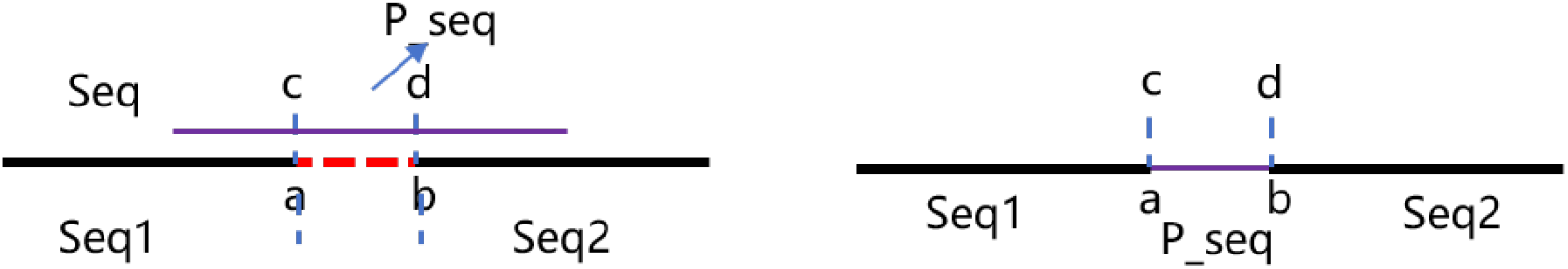
Reading filling processing method

**Fig. 4.**
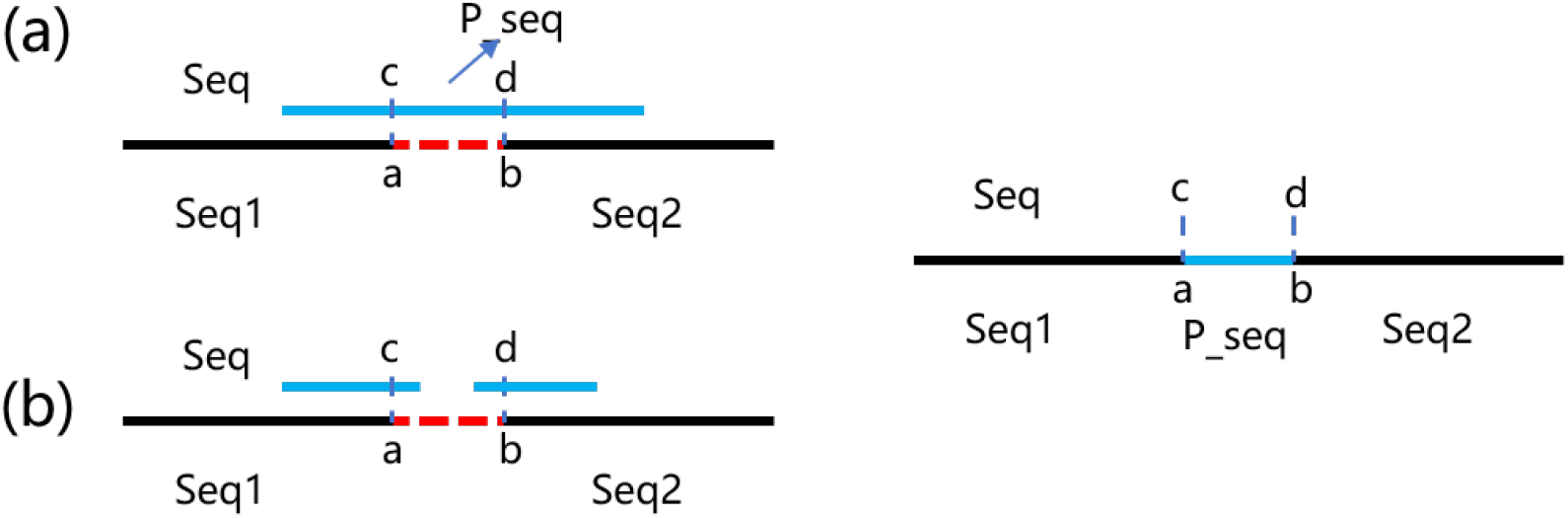
Assembly filling process

**Fig. 5.**
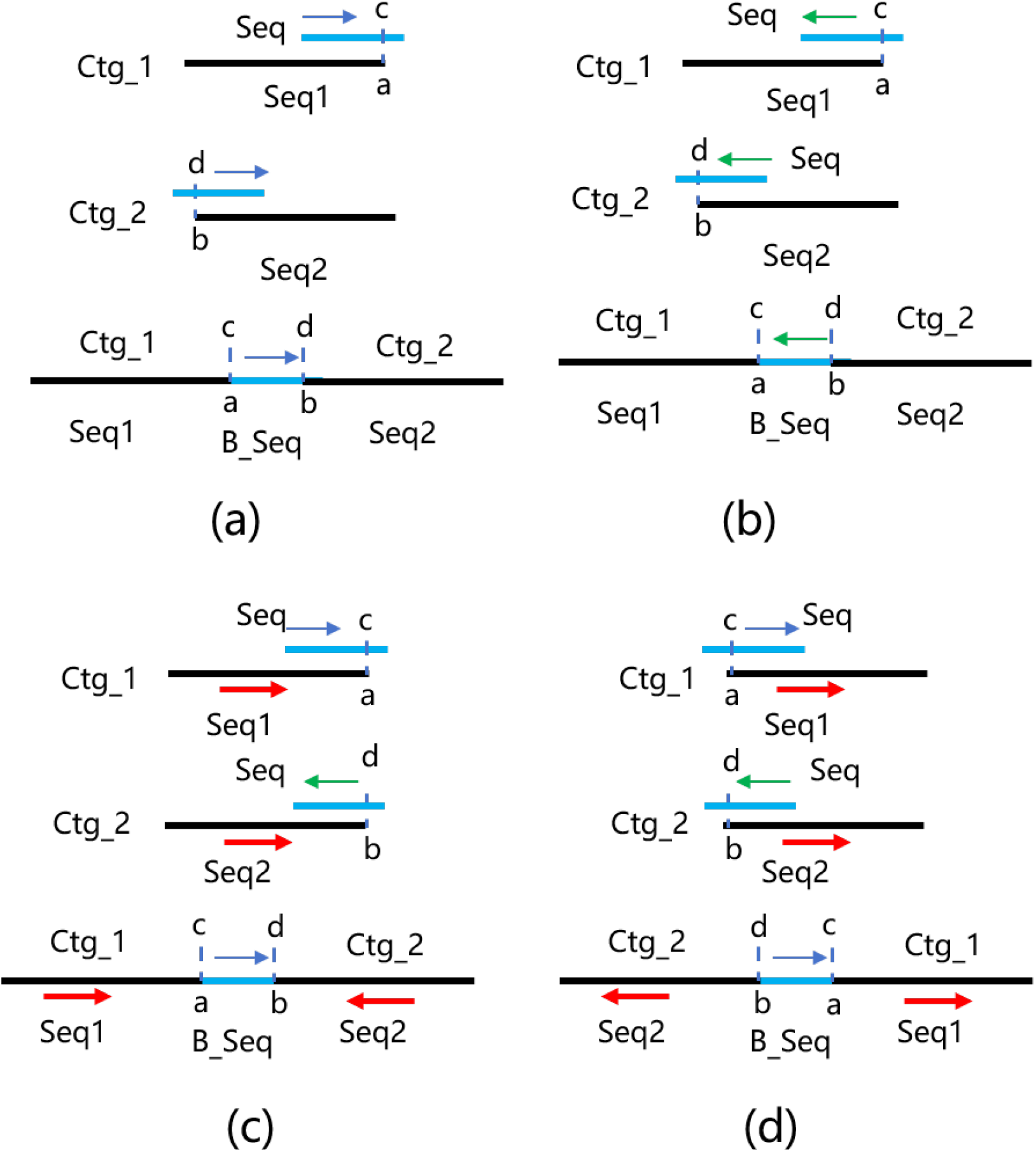
Assembly bridging method

**Fig. 6.**
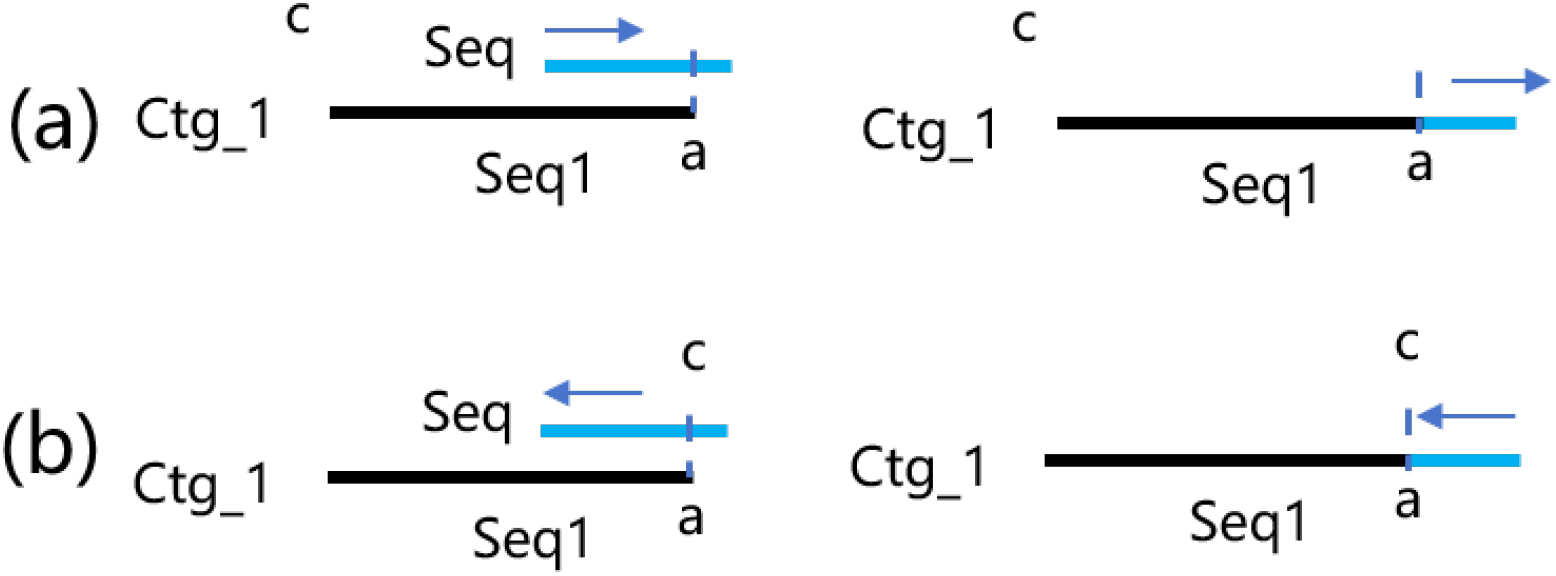
Assembly extension processing method

### 2.6 Polishing

Genome polishing, typically applied after assembly, corrects residual errors like SNPs and indels to improve accuracy. Racon, which uses SIMD-accelerated Partial Order Alignment (POA), offers good scalability and speed. In RefLA’s workflow, previous steps correct larger divergent regions. Remaining small indels and SNPs are addressed by polishing. Specifically, all long reads are re-aligned to the corrected genome, and Racon is used for a final polishing round, which also helps correct minor errors introduced during previous correction steps.

### 2.7 Scaffolding

During divergence correction, RefLA may truncate and bridge sequences. While some sequences are successfully bridged, many may remain unconnected. Under the assumption that long sequence fragments largely maintain their spatial order, reference-based scaffolding tools like RaGOO and Chromosomer have been developed. To leverage the reference genome’s inherent spatial layout, RefLA performs scaffolding on the polished contigs based on their original order in the reference, further improving genome contiguity.

## 3 Results

We evaluated assembler performance using multiple ONT and HiFi sequencing datasets from rice (*Oryza sativa*), green algae (*Chlamydomonas reinhardtii* ), mouse (*Mus musculus*), and human (CHM13). Specific reference genomes for assembly and evaluation were selected. For fairness, some datasets were downsampled to ~40× coverage using rasusa [15] (command: `rasusa --input $in.fq –coverage 40 --genome-size $Size -o $out.fq`). Note: CHM13 used 30*×* ONT and 32*×* HiFi data; *Mus musculus* used 25*×* HiFi data; others were downsampled to 40*×*.

### 3.1 Assembly Quality Evaluation

Tables 1, 2, and Figure 7 present quality assessment results. Note: Mis/Local refers to the count of misassemblies and local misassemblies.

**Table 1.**
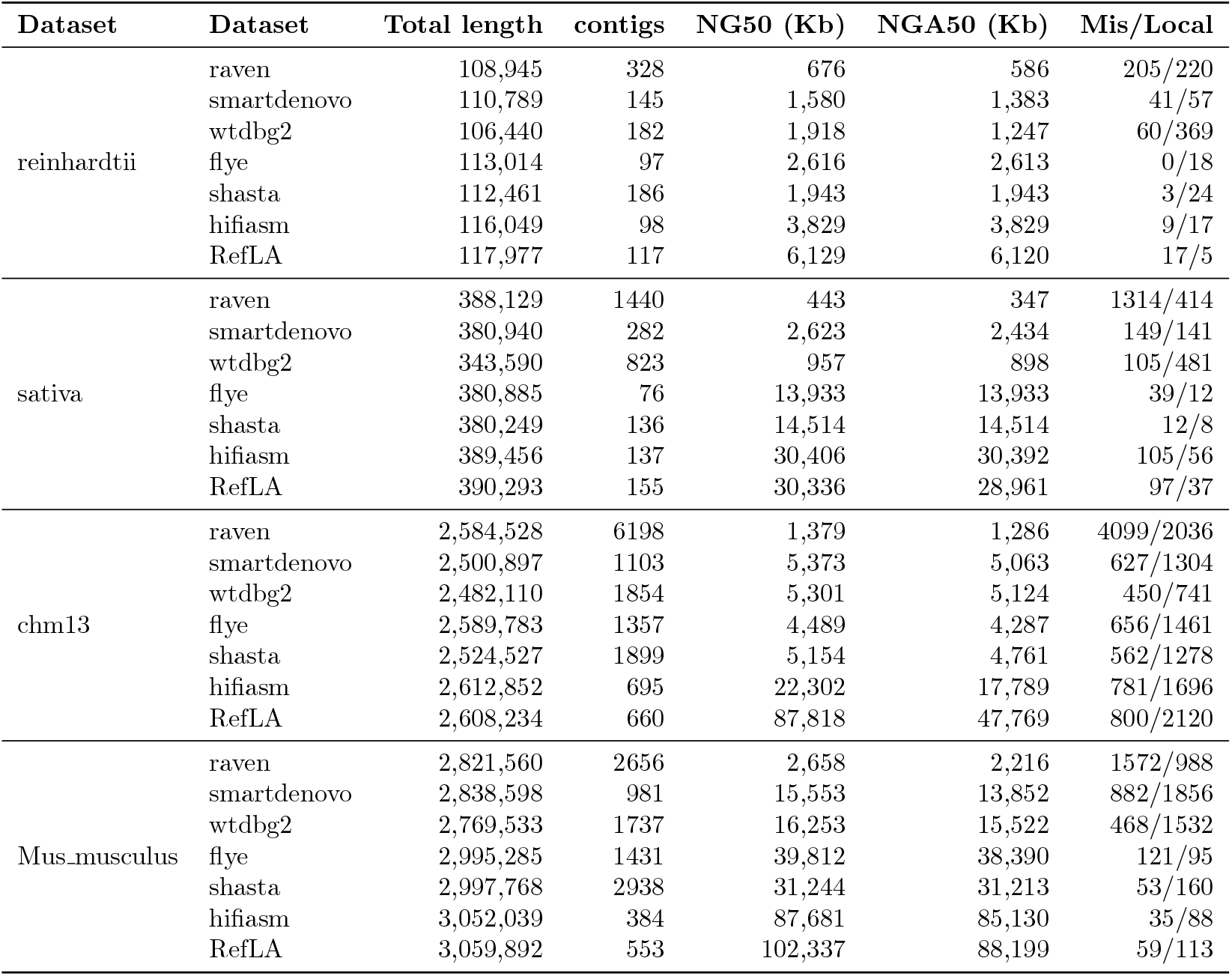
Assembly statistics across datasets and tools. (HiFi data)

| Dataset | Dataset | Total length | contigs | NG50 (Kb) | NGA50 (Kb) | Mis/Local |
| --- | --- | --- | --- | --- | --- | --- |
| reinhardtii | raven | 108,945 | 328 | 676 | 586 | 205/220 |
|  | smartdenovo | 110,789 | 145 | 1,580 | 1,383 | 41/57 |
|  | wtdbg2 | 106,440 | 182 | 1,918 | 1,247 | 60/369 |
|  | flye | 113,014 | 97 | 2,616 | 2,613 | 0/18 |
|  | shasta | 112,461 | 186 | 1,943 | 1,943 | 3/24 |
|  | hifiasm | 116,049 | 98 | 3,829 | 3,829 | 9/17 |
|  | RefLA | 117,977 | 117 | 6,129 | 6,120 | 17/5 |
| sativa | raven | 388,129 | 1440 | 443 | 347 | 1314/414 |
|  | smartdenovo | 380,940 | 282 | 2,623 | 2,434 | 149/141 |
|  | wtdbg2 | 343,590 | 823 | 957 | 898 | 105/481 |
|  | flye | 380,885 | 76 | 13,933 | 13,933 | 39/12 |
|  | shasta | 380,249 | 136 | 14,514 | 14,514 | 12/8 |
|  | hifiasm | 389,456 | 137 | 30,406 | 30,392 | 105/56 |
|  | RefLA | 390,293 | 155 | 30,336 | 28,961 | 97/37 |
| chm13 | raven | 2,584,528 | 6198 | 1,379 | 1,286 | 4099/2036 |
|  | smartdenovo | 2,500,897 | 1103 | 5,373 | 5,063 | 627/1304 |
|  | wtdbg2 | 2,482,110 | 1854 | 5,301 | 5,124 | 450/741 |
|  | flye | 2,589,783 | 1357 | 4,489 | 4,287 | 656/1461 |
|  | shasta | 2,524,527 | 1899 | 5,154 | 4,761 | 562/1278 |
|  | hifiasm | 2,612,852 | 695 | 22,302 | 17,789 | 781/1696 |
|  | RefLA | 2,608,234 | 660 | 87,818 | 47,769 | 800/2120 |
| Mus_musculus | raven | 2,821,560 | 2656 | 2,658 | 2,216 | 1572/988 |
|  | smartdenovo | 2,838,598 | 981 | 15,553 | 13,852 | 882/1856 |
|  | wtdbg2 | 2,769,533 | 1737 | 16,253 | 15,522 | 468/1532 |
|  | flye | 2,995,285 | 1431 | 39,812 | 38,390 | 121/95 |
|  | shasta | 2,997,768 | 2938 | 31,244 | 31,213 | 53/160 |
|  | hifiasm | 3,052,039 | 384 | 87,681 | 85,130 | 35/88 |
|  | RefLA | 3,059,892 | 553 | 102,337 | 88,199 | 59/113 |

**Table 2.** Assembly statistics across datasets and tools. (ONT data)

| Dataset | Dataset | Total length | contigs | NG50 (Kb) | NGA50 (Kb) | Mis/Local |
| --- | --- | --- | --- | --- | --- | --- |
| reinhardtii | raven | 112,686 | 43 | 4,893 | 4,514 | 43/275 |
|  | smartdenovo | 44,868 | 45 | - | - | 3/249 |
|  | wtdbg2 | 112,478 | 447 | 1,621 | 1,167 | 84/993 |
|  | flye | 112,293 | 59 | 5,812 | 5,185 | 7/50 |
|  | shasta | 109,976 | 81 | 5,761 | 4,083 | 24/326 |
|  | RefLA | 112,682 | 76 | 6,747 | 5,772 | 28/313 |
| sativa | raven | 385,393 | 52 | 15,737 | 9,548 | 84/19531 |
|  | smartdenovo | 6,915 | 6 | - | - | 3/346 |
|  | wtdbg2 | 828,891 | 10434 | 564 | 301 | 603/33620 |
|  | flye | 388,439 | 86 | 16,801 | 16,686 | 51/887 |
|  | shasta | 382,196 | 196 | 16,711 | 5,819 | 107/19024 |
|  | RefLA | 388,594 | 160 | 23,122 | 14,851 | 90/3570 |
| chm13 | raven | 2,849,317 | 154 | 46,968 | 45,322 | 255/790 |
|  | smartdenovo | 2,191,811 | 4548 | 389 | 338 | 288/23216 |
|  | wtdbg2 | 3,049,598 | 9389 | 7,128 | 4,498 | 1996/12590 |
|  | flye | 2,856,931 | 618 | 46,584 | 46,584 | 351/845 |
|  | shasta | 2,876,221 | 2182 | 25,347 | 19,799 | 331/3491 |
|  | RefLA | 2,911,708 | 1105 | 57,703 | 48,809 | 351/1515 |

**Fig. 7.**
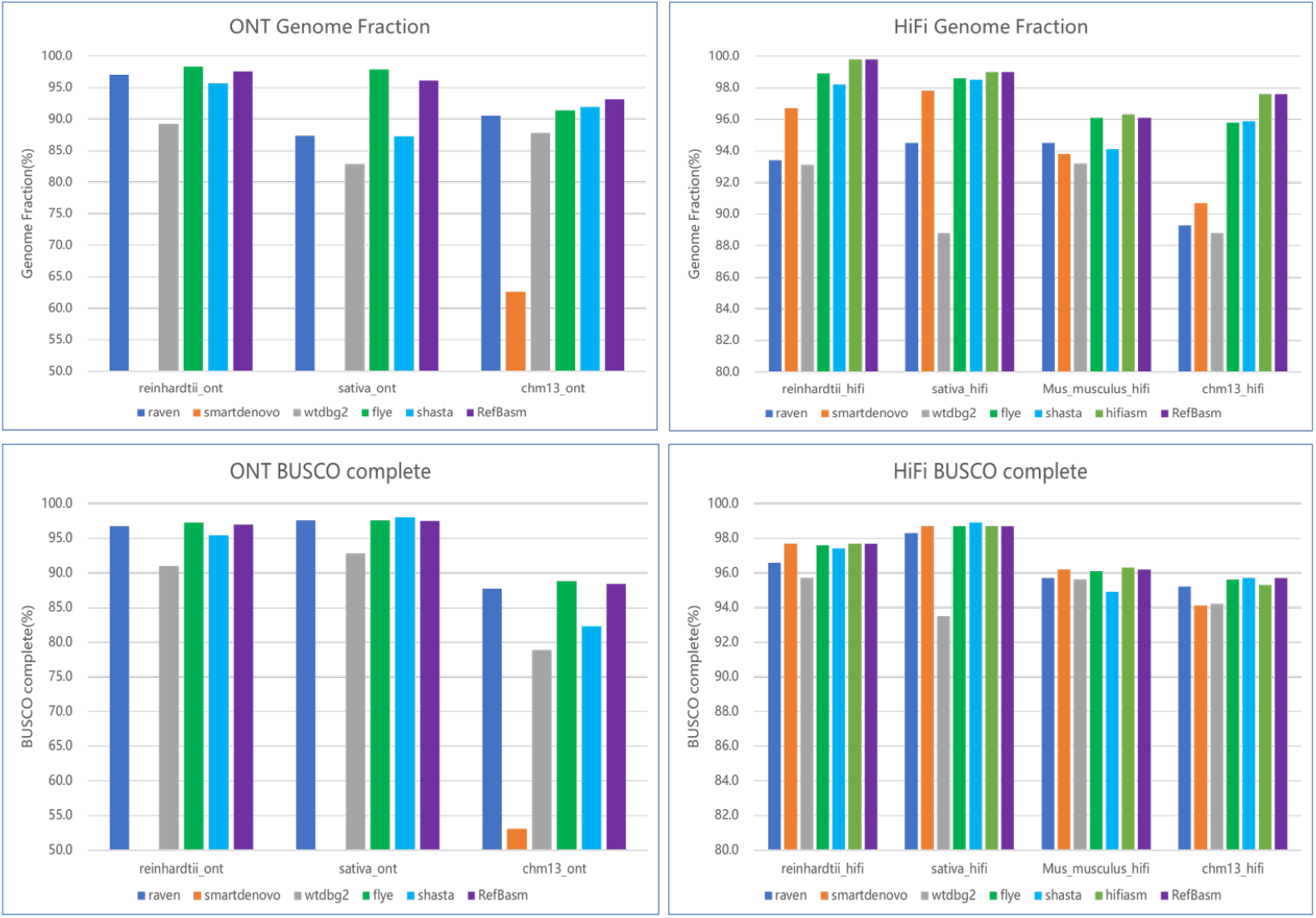
Assembly Results. ONT Data Assembly Integrity Genome Fraction, HiFi Data Assembly Integrity Genome Fraction, ONT data assembly genome integrity score (BUSCO complete score), HiFi data assembly genome integrity score (BUSCO complete score)

Overall, RefLA, Flye, and Hifiasm achieved significantly better assembly quality than other tools. Flye and Hifiasm are currently among the highest-quality de novo assemblers for nanopore and HiFi data, respectively, and are widely used for constructing reference genomes. wtdbg2, Shasta, and Raven are designed for speed, often at the cost of quality. Some tools have specific limitations: Hifiasm is only for HiFi data; Shasta is optimized for human genomes. Considering all datasets and metrics, RefLA, Hifiasm, and Flye generally outperformed other tools, leading in NG50, NGA50, and Genome fraction. Hifiasm excels with HiFi data but is unsuitable for nanopore. Shasta’s performance is less stable on non-human data. SmartDenovo performs poorly on ONT data. In terms of data type adaptability, RefLA and Flye deliver good to excellent results on both ONT and HiFi data. Hifiasm excels on HiFi, Raven performs better on ONT, while wtdbg2 and Shasta show mediocre performance on both.

NG50 and NGA50 reflect assembly contiguity. As shown in Tables 1 and 2, RefLA, Hifiasm, and Flye far surpass wtdbg2 and Shasta, producing more contiguous assemblies. RefLA’s advantage is more pronounced in large genomes. On CHM13 HiFi data, RefLA and Hifiasm achieved N50/NGA50 *≥* 70 Mb, roughly double that of Flye. On CHM13 ONT data, RefLA, Flye, and Raven achieved N50/NGA50 *≥* 40 Mb, approximately double that of Shasta and 8x that of wtdbg2, while SmartDenovo failed to reconstruct the genome.

Genome fraction reflects completeness and the ability to assemble complex, highly repetitive regions. Figure 7 shows that for most species, RefLA, Hifiasm, and Flye report similar, top-tier performance, overall better than wtdbg2, Shasta, Raven, and SmartDenovo. wtdbg2 and Shasta struggle with complex/repeat regions. On CHM13 HiFi and ONT data, RefLA achieved the highest Genome fraction (*≥* 97% and *≥* 93%, respectively), benefiting from the reference genome’s prior information for complex regions. `wtdbg2` and `SmartDenovo` had the lowest Genome fraction (*≤* 90% or failure), indicating their difficulty in assembling complex genomic regions. The human genome contains highly repetitive regions and centromeres; `wtdbg2`’s fuzzy Bruijn graph (FBG) is less effective here.

Assembly correctness is indicated by structural errors (misassemblies/local misassemblies) and base-level quality (QV). Structural error counts relate to N50; assemblers may break contigs at potential errors, leading to shorter N50. As shown in Tables 1 and 2, wtdbg2 and Raven had significantly higher misassembly counts. Hifiasm, Flye, and RefLA better balance contiguity and correctness. For base-level errors (QV), wtdbg2, Shasta, SmartDenovo, and Raven generally lag behind Hifiasm, Flye, and RefLA, especially on the large CHM13 genome. BUSCO scores reflect gene completeness. Figure 7 shows that RefLA, Flye, and Hifiasm achieved the highest BUSCO scores in most datasets, while SmartDenovo and wtdbg2 performed the worst.

In summary, RefLA’s assembly quality on ONT and HiFi data is comparable to high-quality de novo assemblers (Flye, Hifiasm) and superior to fast assemblers (wtdbg2, Shasta, Raven). RefLA is among the best in terms of assembly contiguity.

### 3.2 Assembly Speed Evaluation

Tests were conducted on an HPC platform: standard nodes (Intel Xeon Gold 6248R CPU, 192GB RAM) for small datasets (rice, algae) and fat nodes (Intel Xeon Gold 6248 CPU, 1.5TB RAM) for large genomes (human, mouse). All jobs used 40 threads. Timing results are in Table 3. Overall, Shasta, wtdbg2, Raven, and RefLA were faster than Flye and Hifiasm. SmartDenovo was the slowest. For small/medium genomes, the speed advantage of fast assemblers (wtdbg2, Shasta, Raven) was modest. However, for the large CHM13 genome, RefLA and Shasta were notably faster. This is because RefLA leverages the reference to reduce complexity for large genomes, and Shasta is highly optimized for human data. Notably, wtdbg2 and Raven were not faster than RefLA on large genomes.

**Table 3.** Wall-clock time of assemblers across datasets.

| Tools | reinhardtii_ont | sativa_ont | chm13_ont | Reinhardtii_hifi | Sativa_hifi | Mus_musculu_hiif | chm13_hifi |
| --- | --- | --- | --- | --- | --- | --- | --- |
| raven | 0h11m | 0h47m | 10h28m | 0h8m | 0h46m | 9h14m | 10h32m |
| smartdenovo | 52h25m | 19h15m | 80h10m | 105h52m | 12h5m | 155h48m | 368h34m |
| wtdbg2 | 0h25m | 1h2m | 32h9m | 0h8m | 0h40m | 3h34m | 7h15m |
| flye | 1h39m | 3h8m | 28h12m | 0h39m | 1h52m | 9h37m | 14h32m |
| shasta | 0h5m | 0h16m | 3h4m | 0h29m | 0h54m | 2h10m | 2h55m |
| hifiasm | - | - | - | 0h17m | 1h6m | 3h12m | 5h52m |
| RefLA | 0h12m | 0h39m | 5h1m | 0h16m | 0h33m | 2h32m | 4h50m |

## 4 Conclusions

This paper presents RefLA, a novel reference-guided genome assembly method for rapid, high-quality assembly of new sequencing data using an existing high-quality reference. With an increasing number of species having high-quality reference genomes, reference-guided assembly can accelerate the process and improve quality. RefLA implements a multi-round correction strategy: it first identifies candidate divergent regions from read-to-reference alignments, corrects some regions via read-filling, then performs secondary correction and repair on local assembly regions using three strategies based on local assembly results, and finally applies polishing to fix remaining small-to-medium variants and errors.

We tested RefLA against several state-of-the-art assemblers. Results show that RefLA’s assembly quality metrics are among the best. It excels in assembly contiguity and completeness, particularly for large genomes, benefiting from the reference’s prior information for complex regions. Secondly, RefLA is highly efficient, assembling genomes significantly faster than high-quality assemblers, and comparable to or faster than most fast assemblers, being second only to Shasta in speed. Furthermore, RefLA performs well on both ONT and HiFi data, highlighting the advantages of its reference-guided strategy.

## Declarations

### Ethics approval and consent to participate

Not applicable.

### Consent for publication

Not applicable.

### Code availability

RefLA: is publicly available at GitHub https://github.com/Humonex/RefLA.git

